# Comparison of the predictive values of diagnostic tests subject to a case-control sampling with application to the diagnosis of Human African Trypanosomiasis

**DOI:** 10.1101/638379

**Authors:** José Antonio Roldán-Nofuentes, Saad Bouh Sidaty-Regad

## Abstract

Case-control sampling to compare the accuracy of two binary diagnostic tests is frequent in clinical practice. This type of sampling consists of applying the two diagnostic tests to all of the individuals in a sample of those who have the disease and in another sample of those who do not have the disease. In this sampling, the sensitivities are compared from the case sample applying the McNemar’s test, and the specificities from the control sample. Other parameters of binary tests are the positive and negative predictive values. The predictive values of a diagnostic test represent the clinical accuracy of a binary diagnostic test when it is applied to the individuals in a population with a determined disease prevalence. This article studies the comparison of the predictive values of two diagnostic tests subject to a case-control sampling. A global hypothesis test, based on the chi-square distribution, is proposed to compare the predictive values simultaneously. The comparison of the predictive values is also studied individually. The hypothesis tests studied require knowledge of the disease prevalence. Simulation experiments were carried out to study the type I errors and the powers of the hypothesis tests proposed, as well as to study the effect of a misspecification of the prevalence on the asymptotic behavior of the hypothesis tests and on the estimators of the predictive values. The results obtained were applied to a real example on the diagnosis of the Human African Trypanosomiasis. The model proposed was extended to the situation in which there are more than two diagnostic tests.

## 1. Introduction

The main parameters to assess and compare the accuracy of binary diagnostic tests (BDTs) are sensitivity and specificity. The sensitivity (Se) is the probability of the result of the BDT being positive when the individual has the disease and the specificity (Sp) is the probability of the result of the BDT being negative when the individual does not have the disease. Other parameters that are used to assess and compare two BDTs are the predictive values (PVs). The positive predictive value (PPV) is the probability of an individual having the disease when the result of the BDT is positive, and the negative predictive value (NPV) is the probability of an individual not having the disease when the result of the BDT is negative. The PVs represent the accuracy of the diagnostic test when it is applied to a cohort of individuals, and they are measures of the clinical accuracy of the BDT. The PVs depend on the Se and the Sp of the BDT and on the disease prevalence (p), and are easily calculated applying Bayes’ Theorem i.e.

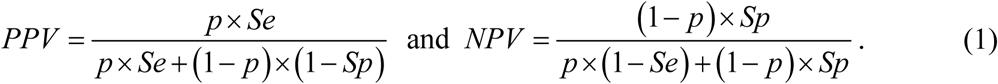

Whereas the Se and the Sp quantify how well the BDT reflects the true disease status (present or absent), the PVs quantify the clinical value of the BDT, since both the individual and the clinician are more interested in knowing how probable it is to have the disease given a BDT result.

The comparison of the performance of two binary diagnostic tests is a topic of special importance in the study of statistical methods for the diagnosis of diseases. This comparison can be made through a cross-sectional sampling or a case-control sampling. Cross-sectional sampling consists of applying the two BDTs and the gold standard to all of the individuals in a single sample. Case-control sampling consists of applying the two BDTs to all of the individuals in two samples, one made up of individuals who have the disease (case sample) and another made up of individuals who do not have the disease (control sample). The advantages and disadvantages of case-control sampling over the cross-sectional can be seen in the book by Pepe (2003). Summarizing, case-control sampling has some advantages over cross-sectional: a) case-control design is more efficient in terms of sample size requirements, b) case-control studies allow for the exploration of subject-related characteristics of the test. Nevertheless, the case-control design has the disadvantage is that by using it we cannot estimate the prevalence of the disease.

The comparison of the sensitivities and the specificities of two BDTs subject to cross-sectional sampling or subject to case-control sampling is made applying the exact comparison test of two paired binomial proportions or McNemar’s test (the asymptotic version of the exact test).

In cross-sectional sampling, the comparison of PVs has been the subject of several studies. Bennett (1972, 1985), Leisenring et al (2000), Wang et al (2006) and Kosinski (2013) studied hypothesis tests to independently compare the PPVs and the NPVs of two BDTs. Moskowitz and Pepe (2006) studied the estimation of the PVs through a confidence region. Roldán-Nofuentes et al (2012) studied the joint comparison of the PPVs and NPVs of two BDTs, and proposed a global hypothesis test based on the chi-square distribution to simultaneously compare the PVs of two BDTs.

In a case-control sampling, Mercaldo et al (2007) have studied the estimation of the PVs of a BDT, assuming that the disease prevalence is known. In this article, we extended the study of Mercaldo et al to the case of two BDTs, studying the comparison of the PVs of the two BDTs subject to a case-control sampling. Subject to a case-control sampling, the two BDTs are applied to all of the individual in two samples, one of *n*_1_ individuals who have the disease (case sample) and another with *n*_2_ individuals who do not have the disease (control sample). In this sampling, the sample sizes *n*_1_ and *n*_0_ are set by the researcher. The sample of individuals that have the disease is extracted from a population of individuals that have the disease (e.g. registers of diseases), and the control sample is extracted from a population of individuals who are known not to have the disease. As the PVs depend on the disease prevalence and subject to a case-control sampling the quotient *n*_1_/(*n*_1_ + *n*_2_) is not an estimator of the prevalence, in order to estimate and compare the PVs subject to this sampling it is necessary to know the prevalence or an estimate of the prevalence. This estimation can be obtained from health surveys or from previous studies. Consequently, the methods of comparison of the PVs subject to a cross-sectional sampling cannot be applied when there is a case-control sampling. In Section 2, we study hypothesis tests to jointly and individually compare the PVs of two BDTs subject to case-control sampling. In Section 3, simulation experiments are carried out to study the type I errors and the powers of the hypothesis tests proposed in Section 2. In Section 4, we study the effect of the misspecification of the prevalence on the asymptotic behavior of the hypothesis tests proposed in Section 2 and on the estimators of the PVs. In Section 5, the results are applied to a real example of the diagnosis of Human African Trypanosomiasis. In Section 6, the model proposed in Section 2 was extended to the situation in which we compare the PVs of more than two BDTs, and in Section 7 the results are discussed.

## 2. The model

Let us consider two BDTs, Test 1 and Test 2, which are applied to all of the individuals in two samples, one of *n*_1_ individuals who have the disease (case sample) and another of *n*_2_ individuals who do not have it (control sample). Let *T*_1_ and *T*_2_ be two binary variables that model the results of each BDT, in such a way that *T*_*i*_ = 1 when the result of the corresponding BDT is positive and *T*_*i*_ = 0 when it is negative. In Table 1, we can see the probabilities associated to the application of both BDTs to both types of individuals (cases and controls), as well as the frequencies observed.

**Table 1.**
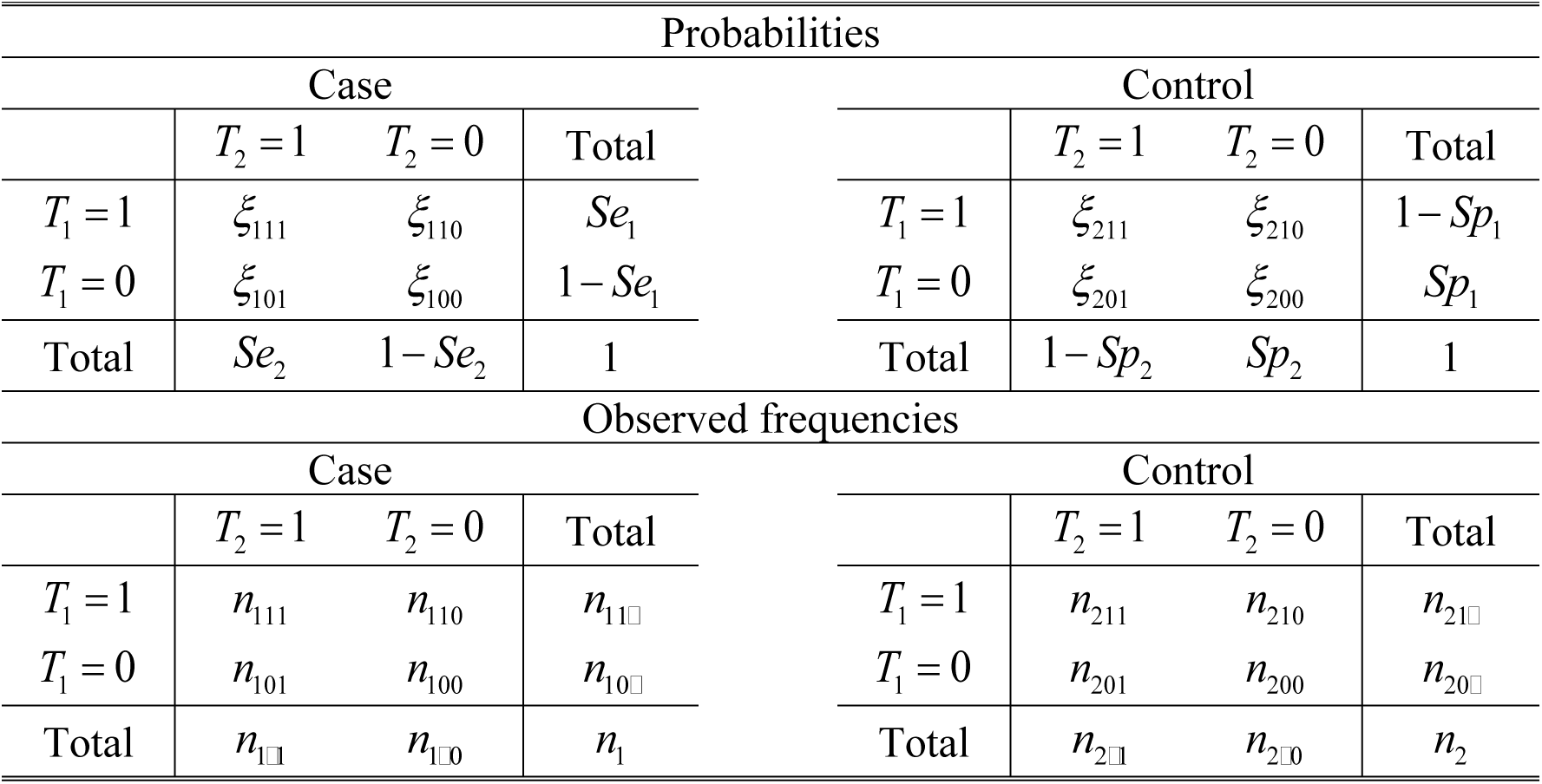
Probabilities and observed frequencies subject to case-control sampling.

Using the conditional dependence model of Vacek (1985), the probabilities given in the table are written as

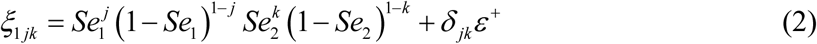

and

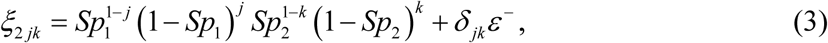

with *j, k* = 0,1. The parameter *ε*^+^ (*ε*^−^) is the covariance between the two BDTs in cases (controls), where *δ*_*jk*_ = 1 if *j* = *k* and *δ*_*jk*_ = −1 if *j* ≠ *k*, and it is verified that 0 ≤ *ε*^+^ ≤ Min{*Se*_1_(1 − *Se*_2_), *Se*_2_(1 − *Se*_1_)} and 0 ≤ *ε*^−^ ≤ Min{*Sp*_1_(1 − *Sp*_2_), *Sp*_2_(1 − *Sp*_1_)}. If *ε*^+^ = *ε*^−^ = 0 then the two BDTs are conditionally independent on the disease status. In practice, the assumption of the conditional independence is not realistic, and therefore *ε*^+^ > 0 and/or *ε*^−^ > 0. In terms of the probabilities *ξ*_*ijk*_, the sensitivities are written as

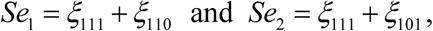

and the specificities as

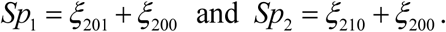

From the case (control) samples *Se*_1_ and *Se*_2_ (*Sp*_1_ and *Sp*_2_) are estimated i.e.

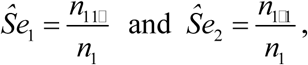

and

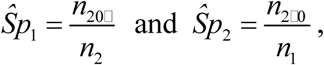

and the estimators of their variances are 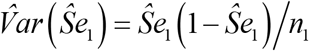, 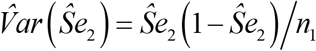, 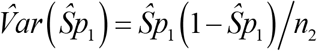 and 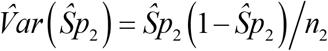. Therefore, the sensitivities and the specificities are estimated as proportions of marginal totals. In this way, in the case sample we are interested in the marginal frequencies *n*_11□_ and *n*_1□1_, and therefore these frequencies are the product of a type I bivariate binomial distribution (Kocherlakota and Kocherlakota, 1992). In an analogous way, from the control sample, the marginal frequencies *n*_20□_ and *n*_2□0_ are the product of a type I bivariate binomial distribution. In the individuals with the disease, the type I bivariate binomial distribution is characterized (Kocherlakota and Kocherlakota, 1992) by the two probabilities *Se*_1_ and *Se*_2_ and by the correlation coefficient (*ρ*^+^) between *T*_1_ and *T*_2_. In an analogous way, in the individuals who do not have the disease, the type I bivariate binomial distribution is characterized by *Sp*_1_, *Sp*_2_ and the correlation coefficient (*ρ*^−^) between *T* and *T*_2_. In the individuals with the disease (cases), the correlation coefficient between the two BDTs is

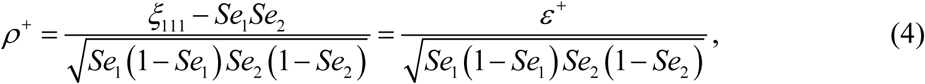

and in the individuals who do not have the disease (controls), the correlation coefficient between the two BDTs is

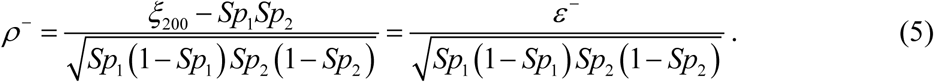

It is easy to check that

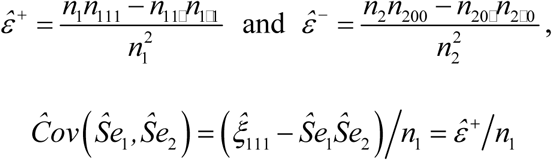

and

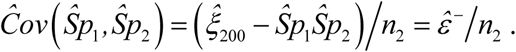

All of the other covariances are zero, since the two samples are independent. The estimators of *ρ*^+^and *ρ*^−^ are

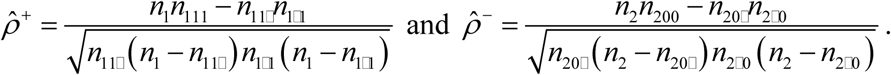

Assuming that prevalence *p* (or an estimation) is known, the estimators of the predictive values are

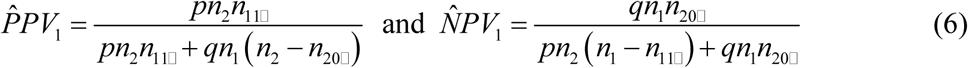

for Test 1, and

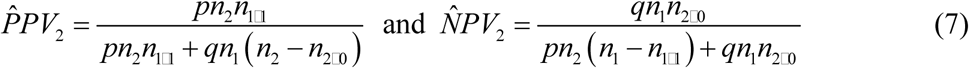

for Test 2, where *q* = 1 − *p*. Let the variance-covariance matrixes be defined as

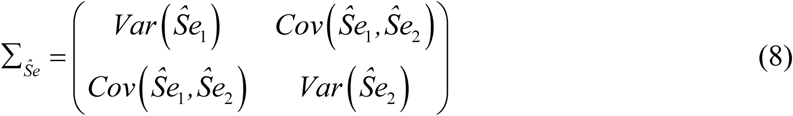

and

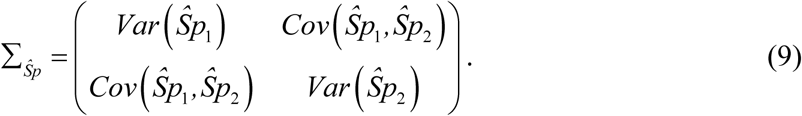

Let **θ** = (*Se*_1_, *Se*_2_, *Sp*_1_, *Sp*_2_)^*T*^ be a vector whose components are the sensitivities and the specificities, and let **ω** = (*PPV*_1_, *PPV*_2_, *NPV*_1_, *NPV*_2_)^*T*^ be a vector whose components are the predictive values. The variance-covariance matrix of 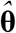 is

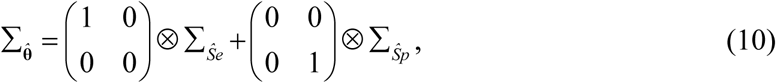

where ⊗ is the product of Kronecker. Applying the delta method, the matrix of variances-covariances of 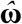 is

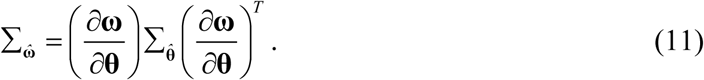

In Appendix A, we can see the expressions of the variances-covariances of the PVs. Then, we study the joint comparison and the individual comparison of the PVs of the two BDTs. In both cases, and as has been explained in Section 1, it is assumed that there is an estimation of the disease prevalence based on a health survey or other studies.

### 2.1. Global hypothesis test

The PVs of each BDT depend on the same parameters, the sensitivity and the specificity of the test and disease prevalence, and therefore they are parameters that depend on each other. Consequently, the PVs of the two BDTs can be compared simultaneously. The global hypothesis test to simultaneously compare the PVs of the two BDTs is

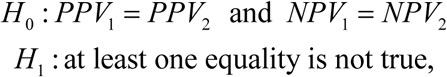

which is equivalent to the hypothesis test

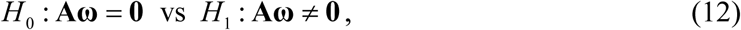

where **A** is a complete range design matrix and a dimension 2 × 4, i.e.

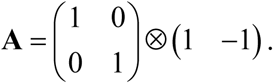

As the vector 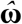 is distributed asymptotically according to a multivariate normal distribution, i.e. 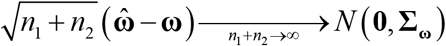, then the statistic for the global hypothesis test (12) is

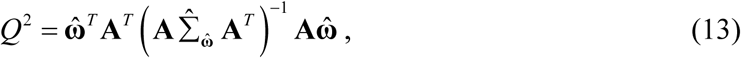

which is distributed asymptotically according to Hotelling’s *T*-squared distribution with a dimension 2 and *n*_1_ + *n*_2_ degrees of freedom, where 2 is the dimension of the vector 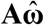. When *n*_1_ + *n*_2_ is large, the statistic *Q*^2^ is distributed according to a central chi-square distribution with 2 degrees of freedom when the null hypothesis is true.

### 2.2. Individual hypothesis tests

The hypothesis test to individually compare the two PPVs (NPVs) is

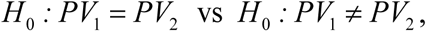

where PV is PPV or NPV. Based on the asymptotic normality of the estimators, the statistic for this hypothesis test is

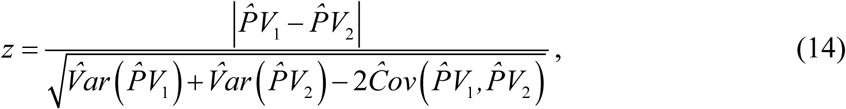

which is distributed asymptotically according to a normal standard distribution, and where the variances-covariances is obtained from the equation (11).

### 2.3. Alternative methods to the global test

The global hypothesis test (12) simultaneously compares the PPVs and the NPVs of the two BDTs. Some alternative methods to this global hypothesis test, based on the individual hypothesis tests, are: 1) Testing the hypotheses *H*_0_ *: PPV*_1_ = *PPV*_2_ and *H*_0_ *: NPV*_1_ = *NPV*_2_ each one to an error *α*; 2) Testing the hypotheses *H*_0_ *: PPV*_1_ = *PPV*_2_ and *H*_0_ *: NPV*_1_ = *NPV*_2_ and applying a multiple comparison method such as Bonferroni’s method (1936) or Holm’s method (1979), which are methods that are very easy to apply based on the p-values. Bonferroni’s method consists of solving each individual hypothesis test to an error *α*/2; and Holm’s method is a step-down method which is based on Bonferroni’s method but is more conservative. In Appendix B, Holm’s method is summarized.

## 3. Simulation experiments

Simulation experiments were carried out to study the type I errors and the powers of the four methods proposed to solve the global hypothesis test: the hypothesis test based on the chi-square (equation (13)), the individual hypothesis tests each one to an error *α*, and the individual hypothesis tests applying Bonferroni’s method and Holm’s method. We have also studied the effect of a misspecification of the prevalence on the asymptotic behaviour of the global hypothesis test and on the estimators of the PVs.

The experiments were designed setting the values of the PVs. For each BDT, we took as PVs the values {0.70, 0.75,…, 0.90, 0.95}, and as disease prevalence we took the values 10%, 25% and 50%. Based on the PVs and the prevalence, the Se and the Sp of each BDT were calculated from the equations (1) and (2), only considering those cases in which the solutions are between 0 and 1. As values of the correlation coefficients *ρ*^+^ and *ρ*^−^ we took low values (25% of the maximum value), intermediate ones (50% of the maximum value) and high ones (75% of the maximum value), and the maximum value of each correlation coefficient is:

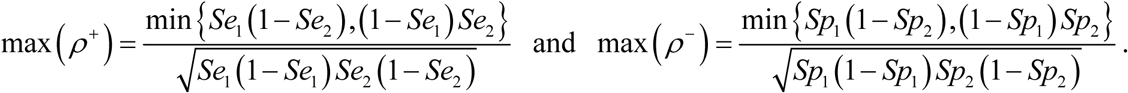

As sample sizes, we took the values *n*_*i*_ = {^50,75,100,200,500^}. The simulation experiments were carried out with R, using the bindata package to generate the samples of each type I bivariate binomial distribution.

Regarding the random samples, these were generated in the following way. Firstly, once the values of the PVs and of the prevalence were set, we calculated the sensitivities and the specificities and maximum values of the coefficients *ρ*^+^ and *ρ*^−^. We then generated 10,000 samples with a type I bivariate binomial distribution with a sample size *n*_1_, probabilities *Se*_1_ and *Se*_2_ and correlation coefficient *ρ*^+^, and another 10,000 samples with a type I bivariate binomial distribution with a sample size *n*_0_, probabilities *Sp*_1_ and *Sp*_2_ and correlation coefficient *ρ*^−^. In this way, we obtained the marginal frequencies *n*_11□_ and *n*_1□1_ (*n*_20□_ and *n*_2□0_) of each one of the 10, 000 case (control) samples. The rest of the marginal frequencies were easily calculated: *n*_10□_ = *n*_1_ − *n*_11□_, *n*_1□0_ = *n*_1_ − *n*_1□1_, *n*_21□_ = *n*_2_ − *n*_20□_ and *n*_2□1_ = *n*_2_ − *n*_2□0_. Then and in order to construct the 2 × 2 table of each case simple, we generated a random valor *n*_111_ of a doubly truncated binomial distribution of parameters *n*_1_ and *ξ*_111_ = *Se*_1_*Se*_2_ + *ε*^+^ *n*_11□_ + *n*_1□1_ − *n*_1_ ≤ *n*_111_ ≤ Min{*n*_11□_, *n*_11□_. This is necessary so that the sum of the frequencies leads to the marginal totals randomly generated through the type I bivariate binomial distribution. In the same way, in order to construct the 2 × 2 table of each control sample, we generated a random value *n*_200_ of a doubly truncated binomial distribution of parameters *n*_2_ and *ξ*_200_ = *Sp*_1_*Sp*_2_ + ε^−^ with *n*_20□_ + *n*_2□0_ − *n*_2_ ≤ *n*_200_ ≤ Min{*n*_20□_,*n*_2□0_. For each one of the 10,000 case (control) samples, once we have generated the values *n*_11□_, *n*_1□1_ and *n*_111_ (*n*_20□_, *n*_2□0_ and *n*_200_) it is easy to construct the complete 2 × 2 table. Thus, *n*_110_ = *n*_1_ − *n*_11□_, *n*_101_ = *n*_1□1_ − *n*_111_ and *n*_100_ = *n*_10□1_ − *n*_101_ for the case samples, and *n*_201_ = *n*_20□_ − *n*_200_, *n*_210_ = *n*_2□0_ − *n*_200_ and *n*_211_ = *n*_21□_ − *n*_210_. For the experiments, the error *α* = 5% was set. Moreover, all of the samples were generated in such a way that in all of them the parameters and the variances-covariances can be estimated.

### 3.1. Type I errors and powers

In Tables 2 and 3, we can see some results obtained for the type I errors of the global test and of the alternative methods proposed in Section 2.3. In these tables, we can only see the results for the global test, the individual comparisons with *α* = 5% and with Bonferroni’s method. The results obtained with Holm’s method are not shown as they are practically the same as those obtained with Bonferroni’s method. From the results obtained we can draw the following conclusions. In general terms, the type I error of the global hypothesis test fluctuates around the error *α* = 5%, especially in the case of samples sized *n*_*i*_ ≥ 100, depending on the prevalence and the correlations between the two BDTs. For samples with smaller sizes (*n*_*i*_ ≤ 75), the type I error of the global test is lower than the error *α* = 5%. The correlations between the two BDTs have an important effect on the type I error of the global test, with a decrease in the type I error when there is an increase in the correlation coefficients. Regarding the method based on the individual hypothesis tests *H*_0_: *PPV*_1_ = *PPV*_2_ and *H*_0_: *NPV*_1_ = *NPV*_2_ to an error *α*= 5% each one of them, the type I error may clearly overwhelm the nominal error (a situation that we have considered when the type I error is greater than 6.5%), especially when the correlations are not high. Consequently, this method may lead to erroneous results (false significances) and, therefore, should not be used. As for solving the global test from the individual tests applying Bonferroni’s (Holm’s) method, the type I error has a very similar behaviour to that of the global hypothesis test.

**Table 2.** Type I errors for *PPV*_1_ = *PPV*_2_ = 0.70and *NPV*_1_ = *NPV*_2_ = 0.95.

|  |  |  |  |  |  |  |  |  |  |  |
| --- | --- | --- | --- | --- | --- | --- | --- | --- | --- | --- |
| $Se_1 = 0.5385, Sp_1 = 0.9744, Se_2 = 0.5385, Sp_2 = 0.9744$ | | | | | | | | | | |
| $0 \leq \rho^+ \leq 1, 0 \leq \rho^- \leq 1$ | | | | | | | | | | |
| $p = 10\%$ | | | | | | | | | | |
| $\rho^+ = 0.25 \quad \rho^- = 0.25$ | | | $\rho^+ = 0.50 \quad \rho^- = 0.50$ | | | $\rho^+ = 0.75 \quad \rho^- = 0.75$ | | | | |
| $n_1$ | $n_2$ | Global | $\alpha = 5\%$ | Bonf. | Global | $\alpha = 5\%$ | Bonf. | Global | $\alpha = 5\%$ | Bonf. |
| 50 | 50 | 0.031 | 0.051 | 0.029 | 0.027 | 0.048 | 0.027 | 0.004 | 0.013 | 0.004 |
| 50 | 75 | 0.029 | 0.059 | 0.029 | 0.025 | 0.051 | 0.026 | 0.004 | 0.017 | 0.005 |
| 50 | 100 | 0.028 | 0.063 | 0.030 | 0.029 | 0.061 | 0.028 | 0.008 | 0.018 | 0.007 |
| 75 | 75 | 0.023 | 0.061 | 0.026 | 0.031 | 0.056 | 0.028 | 0.015 | 0.034 | 0.017 |
| 100 | 100 | 0.027 | 0.063 | 0.029 | 0.023 | 0.052 | 0.024 | 0.020 | 0.043 | 0.019 |
| 200 | 200 | 0.044 | 0.086 | 0.045 | 0.032 | 0.063 | 0.031 | 0.025 | 0.050 | 0.026 |
| 500 | 500 | 0.055 | 0.107 | 0.056 | 0.058 | 0.102 | 0.057 | 0.040 | 0.077 | 0.039 |
| $Se_1 = 0.8615, Sp_1 = 0.8769, Se_2 = 0.8615, Sp_2 = 0.8769$ | | | | | | | | | | |
| $0 \leq \rho^+ \leq 1, 0 \leq \rho^- \leq 1$ | | | | | | | | | | |
| $p = 25\%$ | | | | | | | | | | |
| $\rho^+ = 0.25 \quad \rho^- = 0.25$ | | | $\rho^+ = 0.50 \quad \rho^- = 0.50$ | | | $\rho^+ = 0.75 \quad \rho^- = 0.75$ | | | | |
| $n_1$ | $n_2$ | Global | $\alpha = 5\%$ | Bonf. | Global | $\alpha = 5\%$ | Bonf. | Global | $\alpha = 5\%$ | Bonf. |
| 50 | 50 | 0.048 | 0.094 | 0.046 | 0.018 | 0.047 | 0.018 | 0.001 | 0.007 | 0.002 |
| 50 | 75 | 0.053 | 0.100 | 0.051 | 0.025 | 0.063 | 0.026 | 0.002 | 0.012 | 0.003 |
| 50 | 100 | 0.053 | 0.106 | 0.057 | 0.034 | 0.076 | 0.032 | 0.008 | 0.023 | 0.008 |
| 75 | 75 | 0.059 | 0.105 | 0.055 | 0.039 | 0.087 | 0.037 | 0.007 | 0.016 | 0.006 |
| 100 | 100 | 0.059 | 0.117 | 0.059 | 0.056 | 0.102 | 0.054 | 0.011 | 0.040 | 0.010 |
| 200 | 200 | 0.058 | 0.099 | 0.057 | 0.048 | 0.094 | 0.049 | 0.044 | 0.090 | 0.042 |
| 500 | 500 | 0.052 | 0.098 | 0.053 | 0.051 | 0.101 | 0.052 | 0.049 | 0.090 | 0.048 |
| $Se_1 = 0.9692, Sp_1 = 0.5846, Se_2 = 0.9692, Sp_2 = 0.5846$ | | | | | | | | | | |
| $0 \leq \rho^+ \leq 1, 0 \leq \rho^- \leq 1$ | | | | | | | | | | |
| $p = 50\%$ | | | | | | | | | | |
| $\rho^+ = 0.25 \quad \rho^- = 0.25$ | | | $\rho^+ = 0.50 \quad \rho^- = 0.50$ | | | $\rho^+ = 0.75 \quad \rho^- = 0.75$ | | | | |
| $n_1$ | $n_2$ | Global | $\alpha = 5\%$ | Bonf. | Global | $\alpha = 5\%$ | Bonf. | Global | $\alpha = 5\%$ | Bonf. |
| 50 | 50 | 0.026 | 0.049 | 0.026 | 0.025 | 0.061 | 0.026 | 0.006 | 0.017 | 0.006 |
| 50 | 75 | 0.020 | 0.049 | 0.023 | 0.019 | 0.052 | 0.024 | 0.007 | 0.028 | 0.010 |
| 50 | 100 | 0.019 | 0.043 | 0.023 | 0.016 | 0.045 | 0.019 | 0.010 | 0.034 | 0.014 |
| 75 | 75 | 0.024 | 0.065 | 0.027 | 0.020 | 0.051 | 0.027 | 0.012 | 0.038 | 0.017 |
| 100 | 100 | 0.028 | 0.066 | 0.029 | 0.021 | 0.052 | 0.025 | 0.012 | 0.042 | 0.019 |
| 200 | 200 | 0.047 | 0.088 | 0.044 | 0.034 | 0.074 | 0.032 | 0.021 | 0.058 | 0.026 |
| 500 | 500 | 0.052 | 0.099 | 0.052 | 0.050 | 0.097 | 0.049 | 0.037 | 0.077 | 0.034 |

Global: Global hypothesis test based on the chi-square distribution.
$\alpha = 5\%$ : Individual hypothesis tests each one to an error $\alpha = 5\%$ .
Bonf.: Bonferroni's method.

**Table 3.** Type I errors for *PPV*_1_ = *PPV*_2_ = 0.85 and *NPV*_1_ = *NPV*_2_ = 0.95.

|  |  |  |  |  |  |  |  |  |  |  |
| --- | --- | --- | --- | --- | --- | --- | --- | --- | --- | --- |
| $Se_1 = 0.5312, Sp_1 = 0.9896, Se_2 = 0.5312, Sp_2 = 0.9896$ | | | | | | | | | | |
| $0 \leq \rho^+ \leq 1, 0 \leq \rho^- \leq 1$ | | | | | | | | | | |
| $p = 10\%$ | | | | | | | | | | |
| $\rho^+ = 0.25 \quad \rho^- = 0.25$ | | | $\rho^+ = 0.50 \quad \rho^- = 0.50$ | | | $\rho^+ = 0.75 \quad \rho^- = 0.75$ | | | | |
| $n_1$ | $n_2$ | Global | $\alpha = 5\%$ | Bonf. | Global | $\alpha = 5\%$ | Bonf. | Global | $\alpha = 5\%$ | Bonf. |
| 50 | 50 | 0.033 | 0.056 | 0.034 | 0.020 | 0.051 | 0.024 | 0.004 | 0.014 | 0.004 |
| 50 | 75 | 0.024 | 0.049 | 0.024 | 0.026 | 0.050 | 0.025 | 0.005 | 0.019 | 0.006 |
| 50 | 100 | 0.032 | 0.057 | 0.033 | 0.030 | 0.056 | 0.030 | 0.004 | 0.016 | 0.004 |
| 75 | 75 | 0.034 | 0.054 | 0.033 | 0.025 | 0.052 | 0.026 | 0.014 | 0.036 | 0.015 |
| 100 | 100 | 0.027 | 0.055 | 0.026 | 0.027 | 0.055 | 0.026 | 0.017 | 0.041 | 0.017 |
| 200 | 200 | 0.033 | 0.059 | 0.031 | 0.025 | 0.050 | 0.024 | 0.022 | 0.055 | 0.021 |
| 500 | 500 | 0.046 | 0.087 | 0.049 | 0.031 | 0.068 | 0.033 | 0.018 | 0.050 | 0.024 |
| $Se_1 = 0.85, Sp_1 = 0.95, Se_2 = 0.85, Sp_2 = 0.95$ | | | | | | | | | | |
| $0 \leq \rho^+ \leq 1, 0 \leq \rho^- \leq 1$ | | | | | | | | | | |
| $p = 25\%$ | | | | | | | | | | |
| $\rho^+ = 0.25 \quad \rho^- = 0.25$ | | | $\rho^+ = 0.50 \quad \rho^- = 0.50$ | | | $\rho^+ = 0.75 \quad \rho^- = 0.75$ | | | | |
| $n_1$ | $n_2$ | Global | $\alpha = 5\%$ | Bonf. | Global | $\alpha = 5\%$ | Bonf. | Global | $\alpha = 5\%$ | Bonf. |
| 50 | 50 | 0.023 | 0.058 | 0.022 | 0.005 | 0.030 | 0.007 | 0.001 | 0.005 | 0.001 |
| 50 | 75 | 0.037 | 0.077 | 0.036 | 0.014 | 0.039 | 0.015 | 0.001 | 0.008 | 0.001 |
| 50 | 100 | 0.049 | 0.092 | 0.048 | 0.022 | 0.056 | 0.022 | 0.001 | 0.007 | 0.002 |
| 75 | 75 | 0.042 | 0.087 | 0.041 | 0.025 | 0.055 | 0.025 | 0.004 | 0.014 | 0.004 |
| 100 | 100 | 0.048 | 0.095 | 0.043 | 0.028 | 0.066 | 0.027 | 0.005 | 0.025 | 0.005 |
| 200 | 200 | 0.033 | 0.059 | 0.031 | 0.025 | 0.050 | 0.024 | 0.022 | 0.055 | 0.021 |
| 500 | 500 | 0.048 | 0.097 | 0.046 | 0.056 | 0.101 | 0.051 | 0.050 | 0.099 | 0.049 |
| $Se_1 = 0.9562, Sp_1 = 0.8312, Se_2 = 0.9562, Sp_2 = 0.8312$ | | | | | | | | | | |
| $0 \leq \rho^+ \leq 1, 0 \leq \rho^- \leq 1$ | | | | | | | | | | |
| $p = 50\%$ | | | | | | | | | | |
| $\rho^+ = 0.25 \quad \rho^- = 0.25$ | | | $\rho^+ = 0.50 \quad \rho^- = 0.50$ | | | $\rho^+ = 0.75 \quad \rho^- = 0.75$ | | | | |
| $n_1$ | $n_2$ | Global | $\alpha = 5\%$ | Bonf. | Global | $\alpha = 5\%$ | Bonf. | Global | $\alpha = 5\%$ | Bonf. |
| 50 | 50 | 0.031 | 0.072 | 0.031 | 0.014 | 0.041 | 0.015 | 0.001 | 0.007 | 0.001 |
| 50 | 75 | 0.032 | 0.069 | 0.033 | 0.022 | 0.049 | 0.022 | 0.005 | 0.015 | 0.005 |
| 50 | 100 | 0.025 | 0.057 | 0.026 | 0.025 | 0.064 | 0.026 | 0.008 | 0.025 | 0.008 |
| 75 | 75 | 0.038 | 0.081 | 0.037 | 0.027 | 0.054 | 0.025 | 0.006 | 0.017 | 0.006 |
| 100 | 100 | 0.039 | 0.084 | 0.038 | 0.031 | 0.073 | 0.030 | 0.008 | 0.030 | 0.009 |
| 200 | 200 | 0.033 | 0.059 | 0.031 | 0.025 | 0.050 | 0.024 | 0.022 | 0.055 | 0.021 |
| 500 | 500 | 0.051 | 0.099 | 0.049 | 0.050 | 0.097 | 0.047 | 0.043 | 0.087 | 0.042 |

Global: Global hypothesis test based on the chi-square distribution.
$\alpha = 5\%$ : Individual hypothesis tests each one to an error $\alpha = 5\%$ .
Bonf.: Bonferroni's method.

Regarding the power of the hypothesis tests, in Tables 4 and 5 we can see some of the results obtained for the global test and other alternative methods (Section 2.3). Neither can we see in these Tables the results obtained applying Holm’s method as they are practically the same as those obtained with Bonferroni’s method. From the results, the following conclusions are obtained. The disease prevalence has an important effect on the power of each one of the methods to solve the global test, and the power increases with an increase in the prevalence. Regarding the correlations *ρ*^+^ and *ρ*^−^, these do not have a clear effect on the power, and the power increases sometimes and decreases other times when the correlations increase. In very general terms, when the prevalence is relatively small (*p* = 10%) we need large samples (*n*_i_ > 500) so that the power of the global hypothesis test (equation (13)) is greater than 80%; for a prevalence of 25% with sample sizes *n*_*i*_ ≥ 200 we obtain a power greater than 80%; and for a very large prevalence (*p* = 50%); with sample sizes *n*_*i*_ ≥ 50 we obtain a very higher power, greater than 80%-90%, depending on the difference between the PVs. The power of the method based on the individual hypothesis tests to an error *α* = 5% is greater than that of the global test based on the chi-square distribution due to the fact that its type I error is also greater. Regarding the hypothesis tests based on the individual tests with Bonferroni’s method and Holm’s method, their corresponding power is practically the same, and is also very similar to the power of the global test based on the chi-square distribution.

**Table 4.** Powers for *PPV*_1_ = 0.75, *NPV*_1_ = 0.95, *PPV*_2_ = 0.60 and *NPV*_2_ = 0.95.

| $Se_1 = 0.5357$, $Sp_1 = 0.9802$, $Se_2 = 0.5455$, $Sp_2 = 0.9596$ | | | | | | | | | | |
| --- | --- | --- | --- | --- | --- | --- | --- | --- | --- | --- |
| $0 \leq \rho^+ \leq 0.9805$, $0 \leq \rho^- \leq 0.6933$ | | | | | | | | | | |
| $p = 10\%$ | | | | | | | | | | |
| $\rho^+ = 0.25$ $\rho^- = 0.17$ | | | $\rho^+ = 0.49$ $\rho^- = 0.35$ | | | $\rho^+ = 0.74$ $\rho^- = 0.52$ | | | | |
| $n_1$ | $n_2$ | Global | $\alpha = 5\%$ | Bonf. | Global | $\alpha = 5\%$ | Bonf. | Global | $\alpha = 5\%$ | Bonf. |
| 50 | 50 | 0.025 | 0.056 | 0.031 | 0.023 | 0.049 | 0.024 | 0.005 | 0.019 | 0.007 |
| 50 | 75 | 0.037 | 0.077 | 0.036 | 0.029 | 0.063 | 0.030 | 0.010 | 0.030 | 0.011 |
| 50 | 100 | 0.054 | 0.103 | 0.052 | 0.042 | 0.084 | 0.038 | 0.019 | 0.046 | 0.016 |
| 75 | 75 | 0.038 | 0.078 | 0.038 | 0.032 | 0.066 | 0.033 | 0.018 | 0.042 | 0.018 |
| 100 | 100 | 0.053 | 0.098 | 0.047 | 0.044 | 0.081 | 0.037 | 0.031 | 0.063 | 0.026 |
| 200 | 200 | 0.199 | 0.276 | 0.180 | 0.208 | 0.286 | 0.181 | 0.168 | 0.252 | 0.138 |
| 500 | 500 | 0.495 | 0.575 | 0.462 | 0.591 | 0.668 | 0.556 | 0.720 | 0.785 | 0.678 |
| $Se_1 = 0.8571$, $Sp_1 = 0.9048$, $Se_2 = 0.8727$, $Sp_2 = 0.8061$ | | | | | | | | | | |
| $0 \leq \rho^+ \leq 0.9354$, $0 \leq \rho^- \leq 0.6614$ | | | | | | | | | | |
| $p = 25\%$ | | | | | | | | | | |
| $\rho^+ = 0.23$ $\rho^- = 0.17$ | | | $\rho^+ = 0.47$ $\rho^- = 0.33$ | | | $\rho^+ = 0.70$ $\rho^- = 0.50$ | | | | |
| $n_1$ | $n_2$ | Global | $\alpha = 5\%$ | Bonf. | Global | $\alpha = 5\%$ | Bonf. | Global | $\alpha = 5\%$ | Bonf. |
| 50 | 50 | 0.259 | 0.335 | 0.230 | 0.254 | 0.345 | 0.230 | 0.195 | 0.334 | 0.210 |
| 50 | 75 | 0.409 | 0.496 | 0.378 | 0.454 | 0.543 | 0.424 | 0.467 | 0.606 | 0.470 |
| 50 | 100 | 0.505 | 0.584 | 0.462 | 0.598 | 0.677 | 0.556 | 0.683 | 0.776 | 0.675 |
| 75 | 75 | 0.416 | 0.498 | 0.382 | 0.469 | 0.557 | 0.436 | 0.501 | 0.608 | 0.476 |
| 100 | 100 | 0.528 | 0.606 | 0.488 | 0.625 | 0.699 | 0.579 | 0.718 | 0.793 | 0.685 |
| 200 | 200 | 0.822 | 0.862 | 0.790 | 0.891 | 0.923 | 0.873 | 0.974 | 0.983 | 0.964 |
| 500 | 500 | 0.996 | 0.999 | 0.996 | 1 | 1 | 1 | 1 | 1 | 1 |
| $Se_1 = 0.9643$, $Sp_1 = 0.6786$, $Se_2 = 0.9818$, $Sp_2 = 0.3455$ | | | | | | | | | | |
| $0 \leq \rho^+ \leq 0.7071$, $0 \leq \rho^- \leq 0.5$ | | | | | | | | | | |
| $p = 50\%$ | | | | | | | | | | |
| $\rho^+ = 0.18$ $\rho^- = 0.13$ | | | $\rho^+ = 0.35$ $\rho^- = 0.25$ | | | $\rho^+ = 0.53$ $\rho^- = 0.38$ | | | | |
| $n_1$ | $n_2$ | Global | $\alpha = 5\%$ | Bonf. | Global | $\alpha = 5\%$ | Bonf. | Global | $\alpha = 5\%$ | Bonf. |
| 50 | 50 | 0.890 | 0.939 | 0.893 | 0.935 | 0.969 | 0.941 | 0.977 | 0.989 | 0.978 |
| 50 | 75 | 0.978 | 0.990 | 0.977 | 0.995 | 0.997 | 0.993 | 0.999 | 0.999 | 0.999 |
| 50 | 100 | 0.995 | 0.998 | 0.995 | 0.999 | 0.998 | 0.999 | 1 | 1 | 1 |
| 75 | 75 | 0.984 | 0.992 | 0.983 | 0.995 | 0.999 | 0.994 | 0.999 | 1 | 0.999 |
| 100 | 100 | 0.998 | 0.999 | 0.998 | 1 | 1 | 0.999 | 1 | 1 | 1 |
| 200 | 200 | 1 | 1 | 1 | 1 | 1 | 1 | 1 | 1 | 1 |
| 500 | 500 | 1 | 1 | 1 | 1 | 1 | 1 | 1 | 1 | 1 |

**Table 5.** Powers for *PPV*_1_ = 0.95, *NPV*_1_ = 0.95, *PPV*_2_ = 0.75 and *NPV*_2_ = 0.95.

|  |  |  |  |  |  |  |  |  |  |  |
| --- | --- | --- | --- | --- | --- | --- | --- | --- | --- | --- |
| $Se_1 = 0.5278$, $Sp_1 = 0.9969$, $Se_2 = 0.5357$, $Sp_2 = 0.9802$ | | | | | | | | | | |
| $0 \leq \rho^+ \leq 0.9841$, $0 \leq \rho^- \leq 0.3910$ | | | | | | | | | | |
| $p = 10\%$ | | | | | | | | | | |
| $\rho^+ = 0.25$ $\rho^- = 0.10$ | | | $\rho^+ = 0.49$ $\rho^- = 0.19$ | | | $\rho^+ = 0.74$ $\rho^- = 0.29$ | | | | |
| $n_1$ | $n_2$ | Global | $\alpha = 5\%$ | Bonf. | Global | $\alpha = 5\%$ | Bonf. | Global | $\alpha = 5\%$ | Bonf. |
| 50 | 50 | 0.030 | 0.059 | 0.030 | 0.019 | 0.048 | 0.020 | 0.007 | 0.019 | 0.008 |
| 50 | 75 | 0.031 | 0.063 | 0.032 | 0.023 | 0.054 | 0.023 | 0.009 | 0.024 | 0.009 |
| 50 | 100 | 0.033 | 0.064 | 0.033 | 0.030 | 0.063 | 0.030 | 0.010 | 0.031 | 0.009 |
| 75 | 75 | 0.033 | 0.057 | 0.032 | 0.025 | 0.055 | 0.025 | 0.015 | 0.036 | 0.015 |
| 100 | 100 | 0.034 | 0.067 | 0.033 | 0.026 | 0.059 | 0.026 | 0.026 | 0.054 | 0.026 |
| 200 | 200 | 0.123 | 0.182 | 0.094 | 0.122 | 0.177 | 0.095 | 0.108 | 0.168 | 0.078 |
| 500 | 500 | 0.666 | 0.770 | 0.662 | 0.669 | 0.781 | 0.667 | 0.699 | 0.811 | 0.696 |
| $Se_1 = 0.8444$, $Sp_1 = 0.9852$, $Se_2 = 0.8571$, $Sp_2 = 0.9048$ | | | | | | | | | | |
| $0 \leq \rho^+ \leq 0.9511$, $0 \leq \rho^- \leq 0.3779$ | | | | | | | | | | |
| $p = 25\%$ | | | | | | | | | | |
| $\rho^+ = 0.24$ $\rho^- = 0.09$ | | | $\rho^+ = 0.48$ $\rho^- = 0.19$ | | | $\rho^+ = 0.71$ $\rho^- = 0.28$ | | | | |
| $n_1$ | $n_2$ | Global | $\alpha = 5\%$ | Bonf. | Global | $\alpha = 5\%$ | Bonf. | Global | $\alpha = 5\%$ | Bonf. |
| 50 | 50 | 0.172 | 0.247 | 0.142 | 0.154 | 0.234 | 0.127 | 0.107 | 0.198 | 0.097 |
| 50 | 75 | 0.422 | 0.537 | 0.396 | 0.398 | 0.521 | 0.378 | 0.365 | 0.541 | 0.367 |
| 50 | 100 | 0.627 | 0.734 | 0.615 | 0.641 | 0.755 | 0.638 | 0.674 | 0.779 | 0.653 |
| 75 | 75 | 0.434 | 0.549 | 0.400 | 0.432 | 0.555 | 0.410 | 0.402 | 0.552 | 0.391 |
| 100 | 100 | 0.635 | 0.753 | 0.634 | 0.655 | 0.774 | 0.656 | 0.666 | 0.796 | 0.683 |
| 200 | 200 | 0.965 | 0.981 | 0.964 | 0.977 | 0.987 | 0.974 | 0.989 | 0.994 | 0.988 |
| 500 | 500 | 1 | 1 | 1 | 1 | 1 | 1 | 1 | 1 | 1 |
| $Se_1 = 0.95$, $Sp_1 = 0.95$, $Se_2 = 0.9643$, $Sp_2 = 0.6786$ | | | | | | | | | | |
| $0 \leq \rho^+ \leq 0.8388$, $0 \leq \rho^- \leq 0.3333$ | | | | | | | | | | |
| $p = 50\%$ | | | | | | | | | | |
| $\rho^+ = 0.21$ $\rho^- = 0.08$ | | | $\rho^+ = 0.42$ $\rho^- = 0.17$ | | | $\rho^+ = 0.63$ $\rho^- = 0.25$ | | | | |
| $n_1$ | $n_2$ | Global | $\alpha = 5\%$ | Bonf. | Global | $\alpha = 5\%$ | Bonf. | Global | $\alpha = 5\%$ | Bonf. |
| 50 | 50 | 0.929 | 0.969 | 0.942 | 0.954 | 0.983 | 0.966 | 0.965 | 0.992 | 0.978 |
| 50 | 75 | 0.994 | 0.998 | 0.995 | 0.999 | 0.999 | 0.999 | 0.999 | 1 | 0.999 |
| 50 | 100 | 1 | 1 | 1 | 1 | 1 | 1 | 1 | 1 | 1 |
| 75 | 75 | 0.995 | 0.998 | 0.995 | 0.997 | 0.999 | 0.998 | 1 | 1 | 1 |
| 100 | 100 | 1 | 1 | 1 | 1 | 1 | 1 | 1 | 1 | 1 |
| 200 | 200 | 1 | 1 | 1 | 1 | 1 | 1 | 1 | 1 | 1 |
| 500 | 500 | 1 | 1 | 1 | 1 | 1 | 1 | 1 | 1 | 1 |

As conclusions of the results obtained in the simulation experiments, the global hypothesis test based on the chi-square distribution behaves well in terms of the type I error (it does not overwhelm the nominal error of 5%), the same as the individual tests along with Bonferroni’s method or Holm’s method. The method based on the individual tests to a global error *α* = 5% should not be used as it may clearly overwhelm the nominal error.

From the results obtained, we propose the following method to compare the PVs of two BDTs subject to a case-control sampling: 1) Applying the hypothesis test based on the chi-square distribution (equation (13)) to an error *α*, 2) If the global hypothesis test is not significant, the equality hypothesis of the PVs is not rejected; if the global hypothesis test is significant to an error *α*, the investigation of the causes of the significance is made by testing the individual tests (equation (14)) and applying Bonferroni’s method or Holm’s method to an error *α*.

### 3.2. Effect of the prevalence

The estimation and comparison of the PVs of two BDTs subject to a case-control sampling requires knowledge of the disease prevalence, of an estimation of the disease prevalence obtained from another study, e.g. a health survey. To study the effect of a misspecification of the prevalence in the comparison of the PVs of two BDTs and in the estimators of the PVs, we carried out simulation experiments similar to those made to study the type I errors and the powers. For this purpose, we took as the prevalence for the inference an overestimation (and an underestimation) equal to 5% and to 10% of the value of the prevalence set, and we have studied the type I errors and the powers of the global test and of the Bonferroni and Holm methods and the relative root mean square error (*RRMSE*) of the estimator of each PVs. Thus, for each estimator we calculated the relative root mean square error (RRMSE) defined as

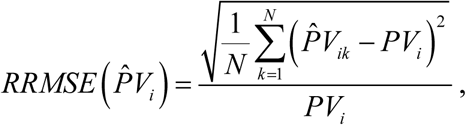

where *PV*_*i*_ is the PPV or the NPV of the *i*th BDT (*i* = 1, 2) and 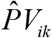 is its estimator calculated from the *k*th sample (*k* = 1,…,*N*), and *N* = 10, 000. For the values of the parameters we took as prevalences *p* ={10%, 25%,50%} respectively, and to estimate the PVs we took as prevalences *p′* = *p* ± *d* × *p* with *d* ={5%,10%}.

In Table 6, we show some of the results obtained for the type I errors and the powers of the global test and the Bonferroni method (the results of the Holms method are not shown as they are practically identically to those obtained with the Bonferroni method). In the Table we show the results when there is no misspecification of the prevalence (*p′* = *p*) and when the prevalence is underestimated (*p′* < *p*) and overestimated (*p′* > *p*). From the results of these experiments, it is verified that the type I errors of the methods studied do not overwhelm the nominal error *α* = 5%, and in general terms there are no important differences between the type I errors when there is a misspecification of the prevalence and when there is not. Regarding the powers, the conclusions are also very similar: there are no important differences between the powers when there is a misspecification of the prevalence and when there is not. Regarding the estimators, in Table 6 we show some of the results obtained for the RRMSEs (in %) of the estimators of the PVs of the two BDTs. There is no important difference between the RRMSEs when there is a misspecification of the prevalence (*p′* < *p* or *p′* > *p*) and the RRMSEs when there is no misspecification of the prevalence (*p′* = *p*). In general terms, this difference is not usually over 5% when the samples are small, and this is even lower when the samples are large. Consequently, misspecifications (5% or 10%) of the disease prevalence do not have any important effect on the type I errors and on the powers of the global hypothesis test and on the alternative methods (Bonferroni and Holm), and nor do they have an important effect on the estimators of the PVs.

**Table 6.** Effect of a misspecification of the prevalence.

| Type I errors |  |  |  |  |  |  |  |  |  |  |  |
| --- | --- | --- | --- | --- | --- | --- | --- | --- | --- | --- | --- |
| $PPV_1 = PPV_2 = 0.90$, $NPV_1 = NPV_2 = 0.80$ | | | | | | | | | | | |
| $Se_1 = 0.2571$, $Sp_1 = 0.9905$, $Se_2 = 0.2571$, $Sp_2 = 0.9905$, $\rho^+ = 0.75$, $\rho^- = 0.75$, $p = 25\%$ | | | | | | | | | | | |
| $n_1$ | $n_2$ | $p' = p = 25\%$ | | $p' = 22.50\%$ | | $p' = 23.75\%$ | | $p' = 26.25\%$ | | $p' = 27.50\%$ | |
|  |  | Global | Bonf. | Global | Bonf. | Global | Bonf. | Global | Bonf. | Global | Bonf. |
| 50 | 50 | 0.001 | 0.001 | 0.001 | 0.002 | 0.001 | 0.001 | 0.001 | 0.001 | 0.001 | 0.001 |
| 50 | 75 | 0.001 | 0.002 | 0.001 | 0.002 | 0.001 | 0.002 | 0.001 | 0.002 | 0.001 | 0.002 |
| 50 | 100 | 0.002 | 0.003 | 0.002 | 0.003 | 0.002 | 0.003 | 0.002 | 0.003 | 0.002 | 0.003 |
| 75 | 75 | 0.003 | 0.007 | 0.003 | 0.008 | 0.003 | 0.008 | 0.003 | 0.007 | 0.003 | 0.007 |
| 100 | 100 | 0.006 | 0.010 | 0.006 | 0.011 | 0.006 | 0.011 | 0.006 | 0.010 | 0.006 | 0.009 |
| 200 | 200 | 0.010 | 0.020 | 0.010 | 0.020 | 0.010 | 0.020 | 0.010 | 0.019 | 0.010 | 0.019 |
| 500 | 500 | 0.020 | 0.024 | 0.020 | 0.024 | 0.020 | 0.024 | 0.020 | 0.024 | 0.020 | 0.023 |
| Powers |  |  |  |  |  |  |  |  |  |  |  |
| $PPV_1 = 0.90$, $PPV_2 = 0.70$, $NPV_1 = 0.80$, $NPV_2 = 0.90$ | | | | | | | | | | | |
| $Se_1 = 0.2571$, $Sp_1 = 0.9905$, $Se_2 = 0.70$, $Sp_2 = 0.90$, $\rho^+ = 0.29$, $\rho^- = 0.22$, $p = 25\%$ | | | | | | | | | | | |
| $n_1$ | $n_2$ | $p' = p = 25\%$ | | $p' = 22.50\%$ | | $p' = 23.75\%$ | | $p' = 26.25\%$ | | $p' = 27.50\%$ | |
|  |  | Global | Bonf. | Global | Bonf. | Global | Bonf. | Global | Bonf. | Global | Bonf. |
| 50 | 50 | 0.999 | 0.992 | 0.999 | 0.993 | 0.999 | 0.992 | 0.999 | 0.992 | 0.999 | 0.991 |
| 50 | 75 | 1 | 0.996 | 1 | 0.995 | 1 | 0.996 | 1 | 0.996 | 1 | 0.995 |
| 50 | 100 | 1 | 0.998 | 1 | 0.997 | 1 | 0.997 | 1 | 0.998 | 1 | 0.998 |
| 75 | 75 | 1 | 1 | 1 | 1 | 1 | 1 | 1 | 1 | 1 | 1 |
| 100 | 100 | 1 | 1 | 1 | 1 | 1 | 1 | 1 | 1 | 1 | 1 |
| 200 | 200 | 1 | 1 | 1 | 1 | 1 | 1 | 1 | 1 | 1 | 1 |
| 500 | 500 | 1 | 1 | 1 | 1 | 1 | 1 | 1 | 1 | 1 | 1 |
| RRMSEs of the estimators of PVs of BDT 1 |  |  |  |  |  |  |  |  |  |  |  |
| $PPV_1 = 0.90$, $PPV_2 = 0.70$, $NPV_1 = 0.80$, $NPV_2 = 0.90$ | | | | | | | | | | | |
| $Se_1 = 0.2571$, $Sp_1 = 0.9905$, $Se_2 = 0.70$, $Sp_2 = 0.90$, $\rho^+ = 0.29$, $\rho^- = 0.22$, $p = 25\%$ | | | | | | | | | | | |
| $n_1$ | $n_2$ | $p' = p = 25\%$ | | $p' = 22.50\%$ | | $p' = 23.75\%$ | | $p' = 26.25\%$ | | $p' = 27.50\%$ | |
| | | $\hat{PPV}_1$ | $\hat{NPV}_1$ | $\hat{PPV}_1$ | $\hat{NPV}_1$ | $\hat{PPV}_1$ | $\hat{NPV}_1$ | $\hat{PPV}_1$ | $\hat{NPV}_1$ | $\hat{PPV}_1$ | $\hat{NPV}_1$ |
| 50 | 50 | 26.8 | 1.8 | 31.1 | 2.7 | 29.3 | 2.1 | 28.3 | 2.1 | 29.7 | 3.2 |
| 50 | 75 | 20.0 | 1.7 | 22.9 | 2.9 | 21.6 | 2.0 | 20.9 | 2.3 | 21.8 | 3.1 |
| 50 | 100 | 15.6 | 1.6 | 18.3 | 3.0 | 16.8 | 1.9 | 15.9 | 2.2 | 17.8 | 3.0 |
| 75 | 75 | 18.4 | 1.4 | 22.3 | 2.5 | 21.2 | 1.7 | 19.3 | 2.0 | 21.9 | 2.8 |
| 100 | 100 | 14.2 | 1.2 | 18.0 | 2.2 | 16.0 | 1.6 | 14.9 | 1.9 | 17.1 | 2.5 |
| 200 | 200 | 8.0 | 0.9 | 9.4 | 1.8 | 8.9 | 1.5 | 8.5 | 1.6 | 8.9 | 1.9 |
| 500 | 500 | 4.1 | 0.5 | 5.4 | 1.5 | 4.6 | 1.2 | 4.5 | 1.2 | 5.1 | 1.6 |
| RRMSEs of the estimators of PVs of BDT 2 |  |  |  |  |  |  |  |  |  |  |  |
| $n_1$ | $n_2$ | $p' = p = 25\%$ | | $p' = 22.50\%$ | | $p' = 23.75\%$ | | $p' = 26.25\%$ | | $p' = 27.50\%$ | |
| | | $\hat{PPV}_2$ | $\hat{NPV}_2$ | $\hat{PPV}_2$ | $\hat{NPV}_2$ | $\hat{PPV}_2$ | $\hat{NPV}_2$ | $\hat{PPV}_2$ | $\hat{NPV}_2$ | $\hat{PPV}_2$ | $\hat{NPV}_2$ |
| 50 | 50 | 10.8 | 2.3 | 14.3 | 2.4 | 13.0 | 2.3 | 11.4 | 2.6 | 13.6 | 2.7 |
| 50 | 75 | 10.3 | 2.2 | 12.4 | 2.2 | 11.1 | 2.2 | 10.8 | 2.5 | 11.8 | 2.6 |
| 50 | 100 | 9.1 | 2.0 | 11.0 | 2.1 | 9.8 | 2.1 | 9.6 | 2.5 | 11.3 | 2.5 |
| 75 | 75 | 10.0 | 1.8 | 12.1 | 2.0 | 10.9 | 1.9 | 10.5 | 2.1 | 12.4 | 2.3 |
| 100 | 100 | 8.9 | 1.6 | 10.6 | 1.8 | 9.5 | 1.7 | 9.3 | 1.9 | 10.9 | 2.1 |
| 200 | 200 | 6.7 | 1.1 | 8.0 | 1.6 | 6.9 | 1.2 | 7.0 | 1.4 | 8.1 | 1.7 |
| 500 | 500 | 4.5 | 0.7 | 5.5 | 1.3 | 4.7 | 0.9 | 4.7 | 1.0 | 5.5 | 1.5 |

Global: Global hypothesis test based on the chi-square distribution.
Bonf.: Bonferroni's method.

## 4. Example

The results obtained were applied to the study by Matovu et al (2010) on the diagnosis of Human African Trypanosomiasis (HAT) in Uganda. HAT, also known as sleeping sickness, is a parasitic disease caused by protozoa belonging to the genus Trypanosoma, and it is transmitted to human beings by a bite from the tsetse fly (genus Glossina) infected by other people or animals that host human pathogenic parasites. In some rural areas of Africa, the disease prevalence may reach 50% in periods of epidemics, and is a significant cause of death. Matovu et al (2010) applied two diagnostic tests to a sample of 75 cases and another sample of 65 controls. In Table 7 (observed frequencies) we can see the frequencies obtained (constructed from the data provided by Matovu et al) when applying the *PCR-Oligochromatography* (*PCR-OC*, variable *T*_1_) test and the *NASBA-Oligochromatography* (*NASBA-OC*, variable *T*_2_) test to both samples of individuals. In order to illustrate the method proposed in this article, two values were considered for the prevalence of HAT: 10% and 50%. The first case (*p* = 10%) corresponds to a situation of low disease prevalence, and the second one (*p* = 50%) corresponds to a situation of a HAT epidemic. In Table 7, we can also see the estimations of the sensitivities and the specificities (and their standard errors, SE) of the BDTs and the correlations.

**Table 7.**
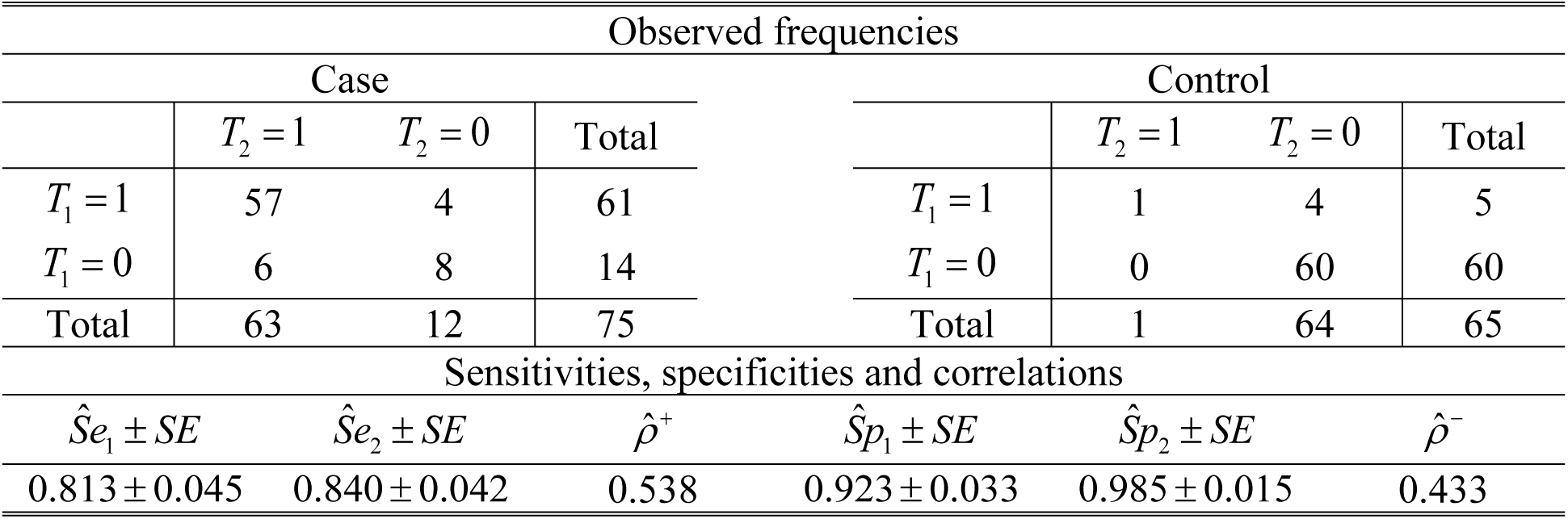
Study by Matuvo et al.

For a prevalence value equal to 10%, the estimations of the PVs are 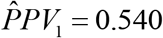, 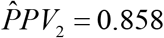, 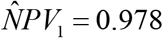 and 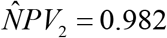, and the estimated variance and covariance matrix of the estimators of the PVs is

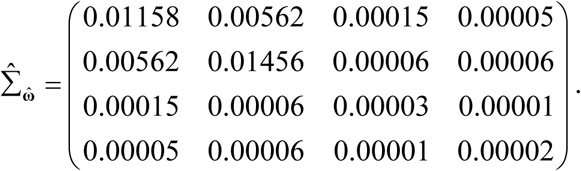

The value of the test statistic for the test

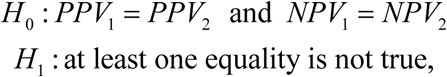

is *Q*^2^ = 6.954 (*P* − *value* = 0.031) and therefore null hypothesis of the global test is rejected. Testing the individual hypothesis tests it is found that the value of the test statistic for the *H*_0_: *PPV*_1_ = *PPV*_2_ is equal to 2.606 (two sided p-value = 0.009), and that the value of the test statistic for the test *H*_0_: *NPV*_1_ = *NPV*_2_ is equal to 0.886 (two sided p-value = 0.375). Applying Bonferroni’s (Holm’s) method the equality hypothesis of the negative predictive values is not rejected and the equality hypothesis of the two positive predictive values is rejected. The positive predictive value of the *NASBA-OC* test is significantly greater than that of the PCR-OC test (95% CI: 0.079 to 0.558).

For a HAT prevalence equal to 50%, the estimations of the PVs are 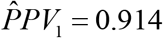, 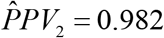, 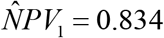 and 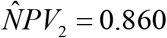, and estimated variance and covariance matrix of the estimators of the PVs is

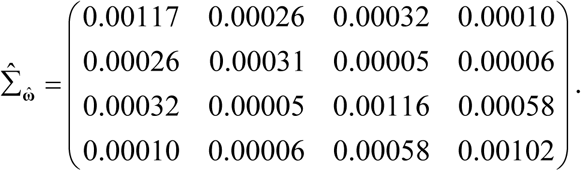

The value of the test statistic for the test

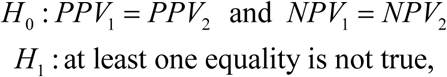

is *Q*^2^ = 5.048 (*P* − *value* = 0.080), and therefore with an error *α* = 5% we do not reject the equality of the positive predictive values and the negative predictive values of both diagnostic tests, although there are signs of significance and, therefore, an increase in the two sample sizes may be necessary. Solving the individual hypothesis tests it is found that the value of the test statistic for the test *H*_0_: *PPV*_1_ = *PPV*_2_ is equal to 2.21 (two sided p-value = 0.027), and that the value of the test statistic for the *H*_0_: *NPV*_1_ = *NPV*_2_ is equal to 0.892 (two sided p-value = 0.373). If each individual hypothesis test is solved to an error *α* = 5%, then *H*_0_: *PPV*_1_ = *PPV*_2_ is rejected and *H*_0_: *NPV*_1_ = *NPV*_2_ is not rejected, and the result is contradictory to that obtained with the global test to an error *α* = 5%.

## 6. More than two BDTs

Let us consider that *J* BDTs (*J* ≥ 3) are applied to all of the individuals in the case sample and the control sample. For each BDT we define the random variable *T*_*j*_ in a similar way to how this was done in Section 2. Let *Se*_*j*_ and *Sp*_*j*_ be the sensitivity and the specificity of the *j*th BDT, with con *j* = 1,…, *J*. Let 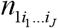 be the number of individuals with the disease for whom *T*_1_ = *i*_1_,…, *T*_*J*_ = *i*_*J*_, with *i*_*j*_ = 1 when the result of the *j*th BDT is positive and *i*_*j*_ = 0 when it is negative. In a similar way, 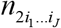 is the number of without the disease for whom *T*_1_ = *i*_1_,…, *T*_*J*_ = *i*_*J*_. Let us consider the probabilities 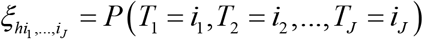, with *h* = 1, 2. Thus, for example for three BDTs, using the dependence model of Torrance-Rynard and Walter (1997), these probabilities are

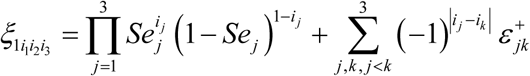

and

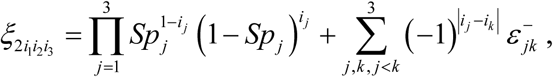

with *i* = 0,1, *i*_*k*_ = 0,1 and *j,k* = 1, 2, 3, and where 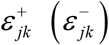 is the covariance between the *j*th BDT and the *k*th BDT for individuals with the disease (without the disease). The estimators of these probabilities are 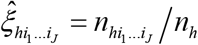, with *h* = 1, 2. The sensitivity and the specificity of the *j*th BDT are

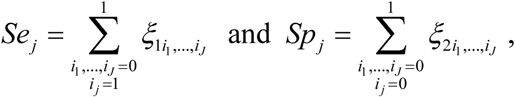

and its estimators are

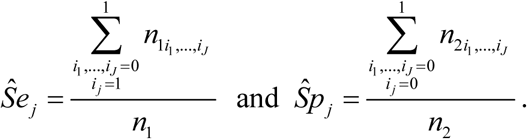

The estimators of the variances-covariances of these estimators are 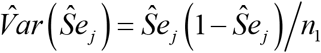, 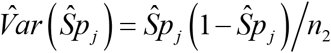, 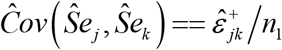 and 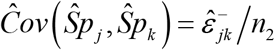, and the rest of the covariances are equal to zero. If we know the prevalence *p* (or and an estimation), the estimators of the PVs of the *j*th BDT are

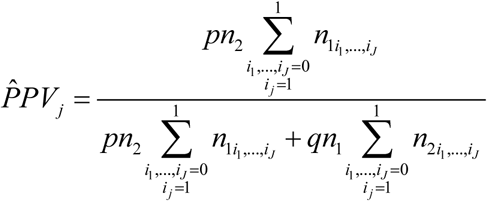

and

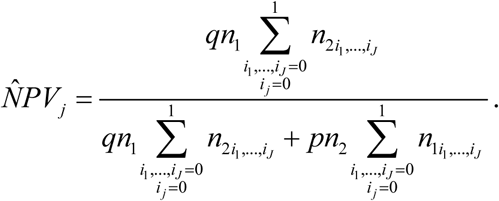

Let **θ** = (*Se*_1_,…, *Se*_*J*_, *Sp*_1_,…, *Sp*_*J*_)^*T*^ be the vector whose components are the sensitivities and the specificities, and let **ω** = (*PPV*_1_,…, *PPV*_*J*_, *NPV*_1_,…, *NPV*_*J*_)^*T*^ be the vector whose components are the PVs. The variance-covariance matrix of 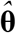, with a dimension 2*J* × 2*J*, is similar to that given in expression (10), where 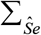 and 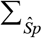 are matrixes with a dimension *J* × *J*. Applying the delta method, the variance-covariance matrix of 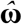, with a dimension 2*J* × 2*J*, has an expression similar to that given in equation (11).

The PVs of each one of the *J* BDTs depend on the same parameters (the sensitivity and the specificity of the *j*th diagnostic test) and, therefore, these parameters can be compared simultaneously. The global hypothesis test to simultaneously compare the PVs of the *J* BDTs is

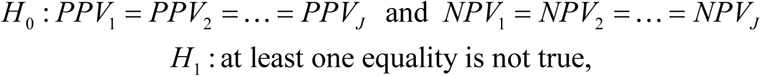

which is equivalent to the hypothesis test

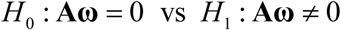

where the matrix **A**, with a dimension 2 (*J* − 1) × 2*J*, is

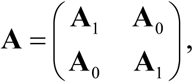

where **A**_0_ is a matrix with a dimension (*J* − 1) × 2*J* whose elements are all equal to 0, and **A**_1_ is a matrix with a dimension (*J* − 1) × 2*J* where each component (*i*, *i*) is equal to 1, each element (*i*, *i* + 1) is equal to −1 for *i* = 1,…, *J* − 1, and the rest of the elements in this matrix are equal to 0. Applying the multivariate central limit theorem it is verified that 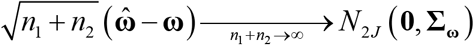. Then, the statistic 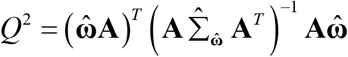 is distributed according to Hotelling’s *T*-squared distribution with a dimension 2(*J* − 1) and *n*_1_ + *n*_2_ degrees of freedom, where 2(*J* − 1) is the dimension of the vector 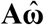. When *n*_1_ + *n*_2_ is large, the statistic *Q*^2^ is distributed according to a central chi-squared distribution with 2(*J* − 1) degrees of freedom when the null hypothesis is true, i.e.

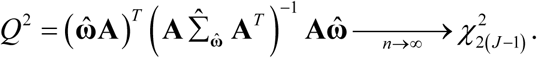

Finally, the method to compare the PVs of the *J* BDTs would consist of the following steps: 1) Solve the global hypothesis test to an error *α*calculating the statistic 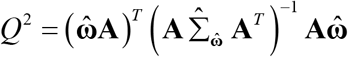 based on the chi-squared distribution; 2) if the global test is not significant to an error *α* then we do not reject the homogeneity of the *J* PVs, but if the hypothesis test is significant then the causes of significance are investigated comparing the PPVs (NPVs) in pairs (equation (14)) and applying an adjustment method of the *p*-value based on multiple comparisons (e.g. Bonferroni or Holm).

## 7. Discussion

The comparison of the positive and negative predictive values of two binary diagnostic tests is an important topic in the study of Statistical Methods in Diagnostic Medicine. Subject to a cross-sectional sampling, this topic has been subject to different studies. In this article we studied the simultaneous comparison of the predictive values of two diagnostic tests subject to a case-control sampling, analysing and comparing several methods. These methods consisted of a global test based on the chi-square distribution, a method based on the individual comparisons each one to a nominal error *α*, and another three methods based on individual comparisons along with a multiple comparison method. The multiple comparison methods that were used were Bonferroni’s method and Holm’s method, which are methods based on the p-values of the individual hypothesis tests and are very easy to apply.

Simulation experiments were carried out to study the type I errors and the power of the four methods proposed. These experiments were based on the generation samples with type I bivariate binomial distributions, which are the distributions that are inherent to case-control design, since from these samples proportions of marginal totals are estimated. The results have shown that the global hypothesis test based on the chi-square distribution behaves well in terms of type I error, and does not overwhelm the nominal error *α* = 5%. Regarding its power, in general this strongly depends on the disease prevalence, and it is necessary to have very large samples (*n*_*i*_ > 500) when the prevalence is small and relatively small sample sizes (*n*_*i*_ > 50) when the prevalence is high, so that the power will be high. The simulation experiments also showed that the methods based on individual hypothesis tests along with multiple comparison methods have type I errors and very similar power to those of the global test based on the chi-square distribution. Consequently, both methods can be used to compare the PVs of the two BDTs. Furthermore, the experiments also showed that the comparison of the predictive values of two diagnostic tests cannot be made independently i.e. comparing the two positive predictive values and comparing the two negative predictive values independently to an error *α* = 5%, as it is possible to obtain a type I error that clearly overwhelms the nominal error set. Based on the results of the simulation experiments, a method has been proposed to compare the predictive values of two diagnostic tests subject to a case-control sampling. This method, which is similar to that proposed by Roldán-Nofuentes et al (2012), consists of the following steps: 1) Simultaneously comparing the predictive values applying the global hypothesis test based on the chi-square distribution (equation (13)) to an error *α*; 2) If the global hypothesis test is not significant, then the equality hypothesis of the PVs is not rejected. If the global hypothesis test is significant to an error *α*, then the causes of the significance are studied solving the individual hypothesis tests (equation (14)) and applying Bonferroni’s method or Holm’s method to an error *α*. This procedure that we propose is similar to the Analysis of Variance: firstly, the global test is solved and, if this is significant, then the causes of the significance are studied starting with paired comparisons along with some multiple comparison method.

Simulation experiments were carried out to study the effect of a misspecification of the prevalence in the asymptotic behaviour of the global hypothesis test based on the chi-square distribution and on the methods based on multiple comparisons. From the results obtained, we can conclude that light or moderate overestimations or underestimations of the prevalence do not have an important effect on the behaviour of these hypothesis tests.

The proposed model has been applied to a real example on the diagnosis of the Human African Trypanosomiasis (HAT) in Uganda, disease that is a major public health problem in some African countries, and whose correct diagnosis is essential for a proper treatment. The results obtained have shown that, when the prevalence is small, the positive predictive value of the NASBA-OC test is significantly greater than that of the PCR-OC test, and there are no significant differences between the negative predictive values of both diagnostic tests. Therefore, when the prevalence of HAT is small, the NASBA-OC test is a better test than the PCR-OC test to confirm the presence of the HAT. When the prevalence of HAT is very high, the equality of the predictive values has not been rejected (although an increase of the two samples may be convenient), and therefore it is not rejected that both diagnostic tests are equally valid to confirm and to exclude the presence of the HAT.

Finally, the global hypothesis test was extended to the situation in which we simultaneously compare the PVs of more than two BDTs, and for this we propose a method which is similar to that proposed for two BDTs. To be able to calculate the global test statistic, 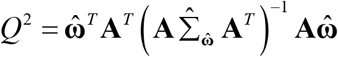, it is necessary for the matrix 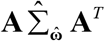 to be non-singular. For two BDTs, the matrix 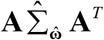 is non-singular when it is verified that *n*_110_ + *n*_101_ > 0 and that *n*_210_ + *n*_201_ > 0; therefore, if *n*_110_ = *n*_101_ = 0 and *n*_210_ = *n*_201_ = 0 then the method proposed to compare the PVs cannot be applied.

## Acknowledgements

This research was supported by the Spanish Ministry of Economy, Grant Number MTM2016-76938-P.

## Appendix A

Performing algebraic operations in equation (11) it is found that:

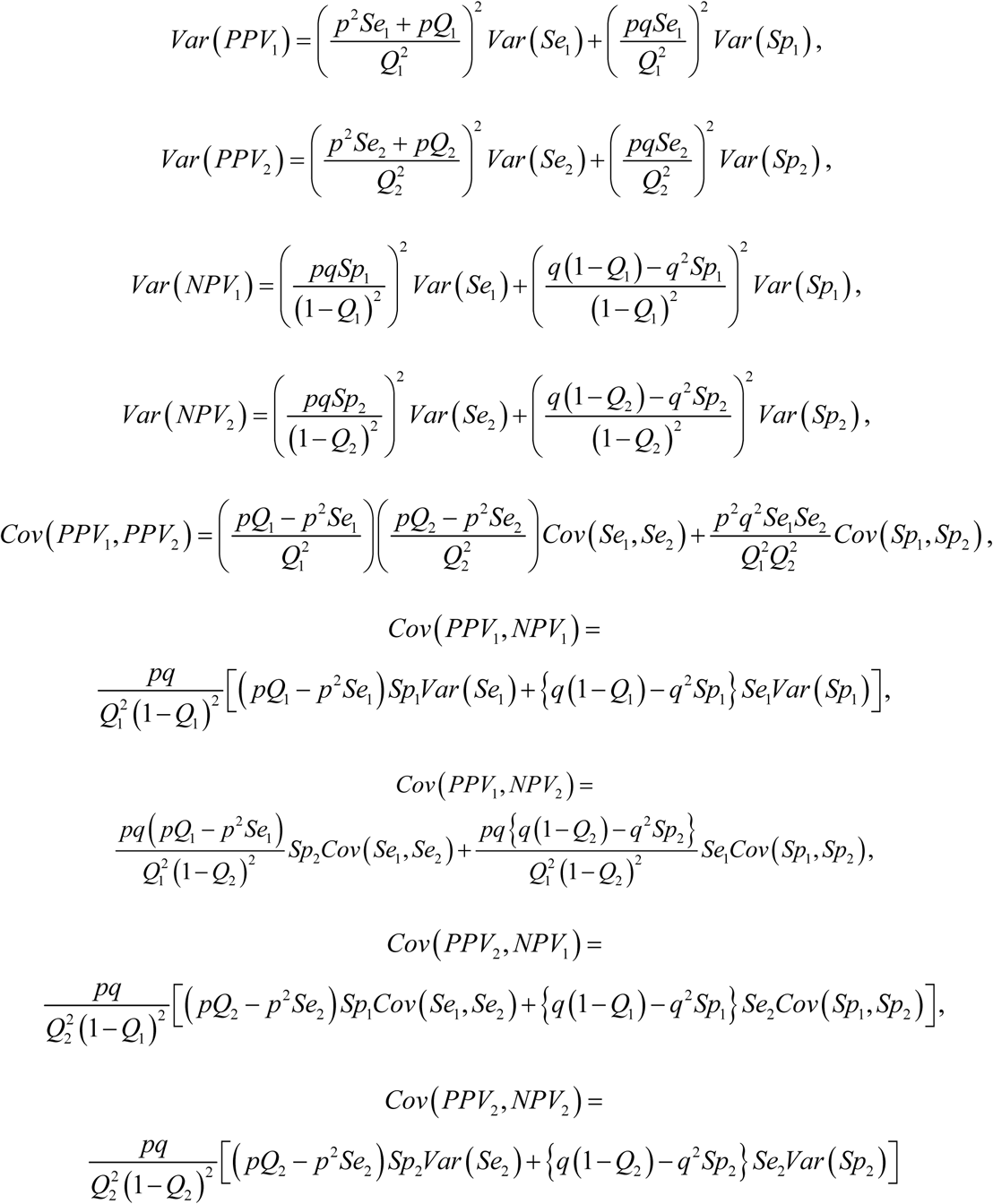

and

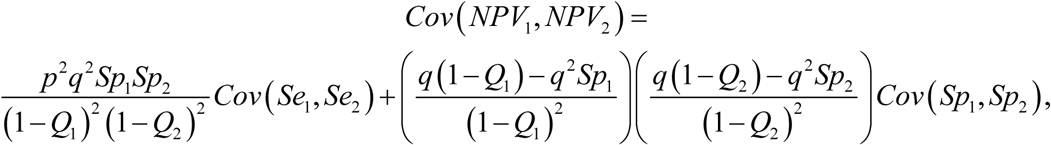

where *q* = 1 − *p* and *Q*_*i*_ = *P* (*T*_*i*_ = 1) = *p* × *Se*_*i*_ + *q* × (1 − *Sp*_*i*_).

## Appendix B

Let us assume that we are going to solve K hypothesis test *H*_0*k*_ vs *H*_1*k*_ with *k* = 1,…, *K*. Let *p*_[1]_ ≤ *p*_[2]_ ≤ … ≤ *p*_[*K*]_ be the *p-values* obtained ordered from the lowest to the highest, and therefore *p*_[*k*]_ is the *p-value* that corresponds to the hypothesis test *H*_0[*k*]_ vs *H*_1[*k*]_. Holm’s method [12] consists of the following steps:

Step 1. If *p*_[1]_ ≤ *α/K* hypothesis *H*_0[*k*]_ is rejected and we go to the next step; if *p*_[1]_ > *α/K* no null hypothesis is rejected and the process finishes.

Step 2. If *p*_[2]_ ≤ *α*/(*K* − 1) hypothesis *H*_0[2]_ is rejected and we go to the next step; if *p*_[2]_ > *α*/(*K* − 1) we do not reject the null hypotheses *H*_0[*k*]_ with *k* = 2,…, *K* and the process finishes….

Step *K*. If *p*_[*K*]_ ≤ *α* hypothesis *H*_0[*K*]_ is rejected and the process finishes; and if *p*_[*K*]_ > *α H*_0[*K*]_ is not rejected and the process finishes.

